# Particle lability drives degradation dynamics and bacterial community assembly during a *Phaeocystis* bloom decline

**DOI:** 10.64898/2026.04.19.716305

**Authors:** Elisa Romanelli, Rebecca Stevens-Green, Carolina Cisternas-Novoa, Julie LaRoche, David A. Siegel, Craig A. Carlson, Uta Passow

## Abstract

Microbial degradation of suspended and sinking organic carbon regulates long-term oceanic carbon storage by controlling the efficiency of the biological pump. Yet microbial controls on carbon export and remineralization remain poorly constrained, limiting predictions of how ocean carbon cycling will respond to climate change. Here, we combined *in situ* sampling with ship-based incubations to quantify prokaryote-driven removal rates of suspended and sinking total organic carbon (TOC). Samples were collected below the mixed layer during three stages of a spring *Phaeocystis pouchetii* bloom in the Labrador Sea. *Phaeocystis* blooms can dominate regional phytoplankton biomass and are expected to increase under future climate. Removal rates were used as a proxy for carbon lability and combined with 16S rRNA metabarcoding and carbon composition analyses to link microbial community structure with substrate characteristics. Removal rates of sinking particles (0.02–0.06 d^−1^) were an order of magnitude higher than those of suspended TOC (0.002 d^−1^) during bloom-decline and non-bloom. In contrast, during late-bloom, suspended carbon exhibited rates of 0.01 d^−1^, comparable to sinking particles, and was enriched in exopolymer-rich colonies. Prokaryotic community composition varied primarily among bloom stages rather than carbon fractions, indicating that bloom stage— and thus particle origin and composition—was the dominant control on bacterial degradation and assembly. Bacterial diversity peaked where carbon was refractory and originated from mixed phytoplankton. Together, these results demonstrate that suspended *Phaeocystis*-derived carbon can be rapidly remineralized when blooms produce exopolymer-rich colonies and highlight bloom stage as key regulator of microbial carbon processing and biological pump efficiency.

## Introduction

Blooms of the cosmopolitan, colony-forming haptophyte *Phaeocystis* contribute substantially to oceanic carbon cycling, attaining surface chlorophyll a concentrations up to 20 mg L^−1^ (Smith et al., 2024). In the Labrador Sea, *Phaeocystis* spp. contributes ∼10% of annual net primary production (Devred et al., 2024), and spring blooms alone can account for ∼20% of seasonal net primary production (present study) (Devred et al., 2025). Given their high productivity, *Phaeocystis* blooms might be expected to contribute substantially to vertical carbon export (Wassmann et al., 1990; Smith et al., 1991). However, most *Phaeocystis* sedimentation events typically only reach upper mesopelagic depths (Reigstad and Wassmann, 2007; Jones and Smith, 2017; Smith et al., 2017; Meyer et al., 2022b), and the contribution of *Phaeocystis*-derived carbon to long-term sequestration is relatively small (Reigstad and Wassmann, 2007; Wolf et al., 2016; Roca-Marti et al., 2025; Laget et al., 2025). Instead, *Phaeocystis*-derived organic matter is thought to be rapidly remineralized by heterotrophic marine bacteria at shallow depths (Beaulieu, 2002). This process may be facilitated by the release of transparent exopolymer particles (TEP) during bloom decline (Mari et al., 2005), which may modify particle composition and enhance bacterial colonization and consumption. However, the microbial mechanisms underlying the limited deep-ocean export of *Phaeocystis*-derived organic matter remain unconstrained.

TEP are acidic polysaccharides retained on a 0.4 μm filter and stained with Alcian Blue (Passow and Alldredge, 1995). During blooms, *Phaeocystis* forms large gelatinous colonies composed of thousands of non-motile cells embedded in a polysaccharide-rich mucilaginous matrix (Hamm et al., 1999; Smith Jr. et al., 2024). Alcian Blue does not stain intact colonies because the colony envelope prevents dye access to the internal matrix (Hamm et al., 1999), and further limits bacterial access to colony-associated carbon and nutrients. Consequently, polysaccharides remain sequestered within intact colonies and become available to heterotrophic bacteria and detectable as TEP only after envelope porosity increases or colonies disintegrate during bloom decline, releasing substantial organic carbon into the surrounding water (Mari et al., 2005). Phytoplankton blooms also produce proteinaceous Coomassie Stainable Particles (CSP); however, little is known about their physical properties (e.g., density, stickiness) and their role in carbon export and sequestration (Cisternas-Novoa et al., 2015).

To assess the susceptibility of *Phaeocystis*-derived organic carbon to microbial degradation across distinct bloom stages, we measured heterotrophic bacterial removal rates of total organic carbon (TOC). TOC removal was quantified for suspended matter and slow- and fast-sinking particles collected using Marine Snow Catchers (MSC, Lampitt et al., 1993) during the decline of a historically large *P. pouchetii* bloom in the eastern Labrador Sea in spring 2022, which extended over more than half of the basin for approximately six weeks (Devred et al., 2025). TOC removal rates were determined from temporal declines in TOC during controlled incubations and used as proxies for organic matter lability. This approach allowed us to examine mechanistic links between the lability of *Phaeocystis*-derived organic matter, including TEP and CSP released during colony formation and decay, and microbial composition across different bloom stages.

## Material and Methods

Data presented in this study were collected in spring 2022 (May 19–June 2) during the Biological Carbon Export in the Labrador Sea (BELAS-1) expedition aboard the R/V *Celtic Explorer*. Distinct stages of *P. pouchetii* bloom termination were sampled, and particulate organic carbon (POC) flux, *P. pouchetii* colony flux, and particle composition were characterized at multiple stations per bloom stage (e.g., Roca-Marti et al., 2025; Laget et al., 2025).

To assess TOC removal rates and microbial community composition, MSCs were deployed below the mixed layer, at the intersection between the primary production zone and the upper mesopelagic at three stations, collecting suspended carbon (dissolved carbon plus suspended particles) and sinking particles. The mixed layer depth was defined as the depth at which potential density differed by 0.03 kg m^−3^ from the surface (Brainerd and Gregg, 1995), and the primary production zone as the depth where fluorescence was 10% of its maximum based on CTD profiles (Roca-Marti et al., 2025). Stations included a late-bloom site (May 22, 150 m; station 9, 58.650° N, 49.360° W), a bloom-decline site (May 27, 130 m; station 28, 57.724° N, 50.066° W), and a non-bloom site (May 30, 100 m; station 30, 59.507° N, 49.674° W). Oceanographic conditions are described in Section 3.1. In addition, water samples were collected using Niskin bottles mounted on a CTD-rosette at six depths between 0 and 500 m at each station to determine TEP, CSP, and particulate organic carbon (POC) concentrations.

### Collection of suspended carbon and sinking particles

Suspended matter and sinking particles were collected using 100-L MSCs with a detachable top that allows separate collection of suspended carbon, slow-sinking, and fast-sinking particles after a predefined settling time of 2 h (Riley et al., 2012; Giering et al., 2016); methodological details are provided in (Romanelli et al., 2023; Romanelli et al., 2024). After settling, three fractions were collected: the top, base, and tray fractions (referred to as MSC fractions). The top fraction, collected from the central tap, contained suspended carbon. To separate slow- and fast-sinking particles, a circular polypropylene tray (diameter 18.5 cm, height 4 cm, volume ∼1 L) was placed at the MSC bottom. After draining the top, the water overlying the tray was collected as the base fraction, while particles retained in the tray constituted the tray fraction. Concentrations of TOC, POC, and microbial community composition based on *16S* rRNA gene sequencing were determined for all three fractions.

TOC and POC concentrations were calculated such that suspended carbon corresponded to the top fraction, while slow- and fast-sinking particle concentrations were derived by subtracting the top fraction from the base fraction and the base fraction from the tray fraction, respectively. These differences were scaled to account for volume ratios (V_MSC = 89.8 L, V_base = 5 L, V_tray = 1 L; h_MSC = 1.5 m). Based on the MSC height and a 2-h settling time, the maximum sinking velocity of slow-sinking particles was 18 m d^−1^ (Giering et al., 2016). The sinking velocity of fast-sinking particles are not further constrained by the MSC. Hereafter, suspended carbon, slow-sinking particles, and fast-sinking particles are collectively referred to as carbon fractions.

### Experiments to measure total organic carbon removed by bacteria

Concentrations and removal rates of TOC were determined from dark-incubation experiments (duration: 11 days) conducted on unfiltered MSC top, base, and tray fractions. Incubations were performed at 4.4 °C in the dark using a refrigerated incubator, within 1 °C of *in situ* temperatures. Because microbial respiration is highly temperature sensitive (Hoppe et al., 2008; Iversen and Ploug, 2013), maintaining near–*in situ* conditions ensured comparability across experiments. Each fraction was incubated in 15 x 40 mL pre-combusted (450 °C, 4 h) borosilicate glass vials (triplicates at five time points). No visible zooplankton was present; although protist contributions cannot be excluded, heterotrophic activity was assumed to be dominated by bacteria (Cohn et al., 2024). At each time point, triplicate samples were fixed with 50 µL DOC-free 4 N HCl (final pH ∼2–3), terminating microbial activity and preventing further TOC remineralization.

TOC concentrations were measured by high-temperature combustion using a modified Shimadzu TOC-V analyzer (Halewood et al., 2022), with analytical precision within ±0.7 µM C. All TOC concentrations exceeded 50 µM C. TOC concentrations for suspended, slow-sinking, and fast-sinking fractions were derived from TOC measurements. Temporal trends in TOC within each carbon fraction were assessed by weighted linear regression of mean values across time points, with standard deviations from three technical replicates. Regressions were performed in MATLAB using fitlm with inverse-variance weighting (Weights = 1./SD.^2), yielding slope, intercept, R^2^, and p-values. Differences among temporal slopes were tested using analysis of covariance (ANCOVA) with time as the covariate. Specific TOC removal rates (d^−1^) were calculated as the slope of the regression of TOC concentrations normalized to Day 0 over the full incubation period (**Table S1**). TOC associated with slow-sinking particles at Day 0 at the late-bloom and bloom-decline stations was excluded due to anomalously low values of unknown origin.

### Measurements of Particulate Organic Carbon (POC)

Concentrations of POC were determined by first filtering duplicate subsamples from the three MSC fractions and water samples collected with Niskin bottles onto pre-combusted (450 °C, 5 hours) GF/F filters (25 mm, Whatman, UK). Filters were dried at 40°C for 12 hours and stored at room temperature. After fuming with HCl for 12 hours, samples were analyzed on a Perkin Elmer 2400 CHN Analyzer with acetanilide standards. The accepted precision of elemental analysis results is 0.4%. Before calculating the POC concentrations, the measured mass was blank corrected (Graff et al., 2023). Blanks (n = 2), prepared by rinsing filters with 0.2-μm filtered seawater, yielded 2.3 ± 0.08 μg POC. Samples yielded values greater than the detection limit and the blank.

### Transparent Exopolymer Particles (TEP) and Coomassie Stainable Particles (CSP)

Samples for TEP and CSP analyses were collected using a CTD/Niskin rosette at station 9 (late-bloom; 10 depths, 5–2000 m) and station 28 (bloom-decline; 11 depths, 5–3500 m). Subsamples (0.5–2 L) from each Niskin bottle were filtered. TEP and CSP were quantified colorimetrically following Passow and Alldredge (1995) and Cisternas-Novoa et al. (2014), with TEP calibration curve prepared following Bittar et al. (2018). Concentrations were normalized to POC to account for differences in particle abundance and are reported as µg XG_eq µg^−1^ POC for TEP and µg BSA_eq µg^−1^ POC for CSP.

In addition, TEP and CSP were analyzed microscopically in the suspended fraction at station 6 (late-bloom; May 21; 58.8° N, 46.4° W), sampled one day prior to station 9 (Alldredge et al., 1993; Long and Azam 1996). The suspended material collected at 80 m was filtered onto 0.4 µm polycarbonate filters and stained onboard with Alcian Blue (TEP) or Coomassie Brilliant Blue (CSP). Filters were mounted on the white side of semitransparent glass slides (CytoClear, Poretics Corp., Livermore, USA) and stored at −20 °C until analysis (Logan et al., 1994). Images were acquired at 200× magnification using a light microscope (Olympus CX41RF) equipped with a digital camera (Lumenera 2-1R, 1.4 MP CCD).

### Assessment of microbial community composition of MSC fractions

Between 200 and 1000 mL of samples from the three MSC fractions (top, base, and tray) were vacuum-filtered onto 25 mm, 0.2 µm polycarbonate Nuclepore filters and stored at −80 °C. DNA was extracted using the DNeasy Plant Mini Kit (Qiagen, Germany) with modifications to the lysis step, including a 5 min incubation with 50 µL of 5 mg mL^−1^ lysozyme (Fisher BioReagents, UK) followed by a 1 h incubation with Buffer AP1 and 45 µL of 20 mg mL^−1^ Proteinase K (Fisher BioReagents, UK) (Zorz et al., 2019). DNA was eluted twice in 50 µL AE buffer and quantified using a NanoDrop 2000 (Thermo Scientific, USA). The V4–V5 region of the *16S* rRNA gene was sequenced using Illumina MiSeq (2 × 300 bp) at the Integrated Microbiome Resource (IMR; Dalhousie University). Libraries were prepared following Comeau et al. (2017) using primers 515FB (GTGYCAGCMGCCGCGGTAA) and 926R (CCGYCAATTYMTTTRAGTTT) (Parada et al., 2016; Walters et al., 2016).

Sequencing data were processed using the Fuhrman lab pipeline in QIIME2 (v2022.2) with DADA2 (v1.22.0), optimized for the 515Y/926R primer set (McNichol et al., 2021; Yeh et al., 2021; Callahan et al., 2016; Liu et al., 2023). Reads were trimmed with cutadapt, separated in silico into 16S and 18S using the SILVA and PR2 databases, and dereplicated, merged, and denoised using the DADA2 consensus method. The forward and reverse 16S reads were truncated at 220 bp and 180 bp, respectively. Taxonomy was assigned using the SILVA database (v138.1), with chloroplast and mitochondrial sequences further classified using PR2 PhytoRef (v4.14.0) (Quast et al., 2013; Yilmaz et al., 2014; Decelle et al., 2015).

Microbiome analyses were conducted in R (v4.5.0) using phyloseq (v1.52.0) and microbiome (v1.30.0). Alpha diversity was calculated using the Shannon index after rarefaction with the ‘rarefy_even_depth’ function. Beta diversity was assessed using principal component analysis (PCA) based on Aitchison distance (Gloor et al., 2017), and differences among stations and MSC fractions were tested using PERMANOVA. Indicator species analysis was performed on CLR-transformed amplicon sequence variant (ASV) data using the multipatt function in the indicspecies package, with bloom stage and carbon fraction treated as separate conditions (De Cáceres and Legendre, 2009). Statistical significance was evaluated at α = 0.05.

## Results

### Oceanographic characterization of P. pouchetii bloom stages

Sampling three stations spanning distinct stages of *P. pouchetii* bloom termination allowed us to place microbial TOC removal rates within an ecological framework. Underwater Vision Profiler (UVP) [44] imagery showed that at MSC deployment depths (112–175 m), intact *P. pouchetii* colonies (0.6–3 mm) were > fivefold more abundant at the late-bloom station than at the bloom-decline and non-bloom stations (Laget et al., 2025). Because senescent *P. pouchetii* colonies fragment (Mari et al., 2025), high abundances of intact colonies suggest a dominance of fresh material. Pigment biomarkers and net primary production data further support this interpretation: the late-bloom station exhibited the highest water-column Chl *a* concentration and the lowest pheopigment-to-Chl *a* ratios (**Figure S1**), indicative of more freshly produced, labile organic matter (Sun et al., 1993). In addition, integrated net primary production was fivefold higher at the late-bloom station than at the bloom-decline stations (20–25 May) (Roca-Marti et al., 2025). Together, these observations indicate that fresh, labile organic matter was available below the mixed layer at the late-bloom station.

At the bloom-decline station, Chl *a* concentrations at deployment depth were twofold lower and the degradation ratios (pheopigment-to-Chl *a*) were fourfold higher than at the late-bloom station (**Figure S1**), consistent with a shift toward more degraded organic matter. Similar transitions have been observed in mesocosm studies of *Phaeocystis globosa*, where increasingly refractory products accumulated as colonies decayed (Brussaard et al., 2005; Alderkamp et al., 2007).

The non-bloom station represented the most oligotrophic conditions, with Chl *a* concentrations an order of magnitude lower and degradation ratios an order of magnitude higher than at the late-bloom station (**Figure S1**). An elevated degradation ratio indicates that water-column carbon, primarily suspended, was dominated by refractory detritus, which is less accessible to microbial degradation (Baltar et al., 2021). Surface NO_3_ concentrations were at or below detection limits, and SiO_4_ in the primary production zone was reduced by ∼50% relative to the other stations (**Figure S1**), suggesting NO_3_ limitation and recent or ongoing silicifier activity. Consistent with nutrient concentrations, diatoms (Stramenopiles) accounted for a relatively high abundance between 5 and 160 m (6–28%) at the non-bloom station compared to the other stations (3–17%; Stevens-Green, *pers. comm*.). Together, these observations indicate a non-bloom system dominated by refractory detritus, likely derived in part from diatom carbon.

### Initial concentrations of POC and TOC of suspended and sinking particles

Mean (± s.d.) POC concentrations of suspended particles were highest at the late-bloom station (6.0 ± 0.4 µmol C L^−1^, 150 m) and lowest at the non-bloom station (3.8 ± 0.3 µmol C L^−1^, 90 m) (**Figure 1A**). Among sinking particles, the highest POC concentrations in slow-sinking particles occurred at the late-bloom station (0.6 ± 0.3 µmol C L^−1^) and in fast-sinking particles at the bloom-decline station (0.8 ± 0.1 µmol C L^−1^) (**Figure 1A**). TOC concentration of suspended carbon (average: 57.6 µmol C L^−1^) was highest at the late-bloom station and lowest at the bloom-decline station and was ∼9-fold greater than the POC concentrations across all stations (**Figure 1B, Figure 2A**). A dominance of dissolved rather than particulate carbon to TOC is common in the ocean (Hansell et al., 2009).

**Figure 1.**
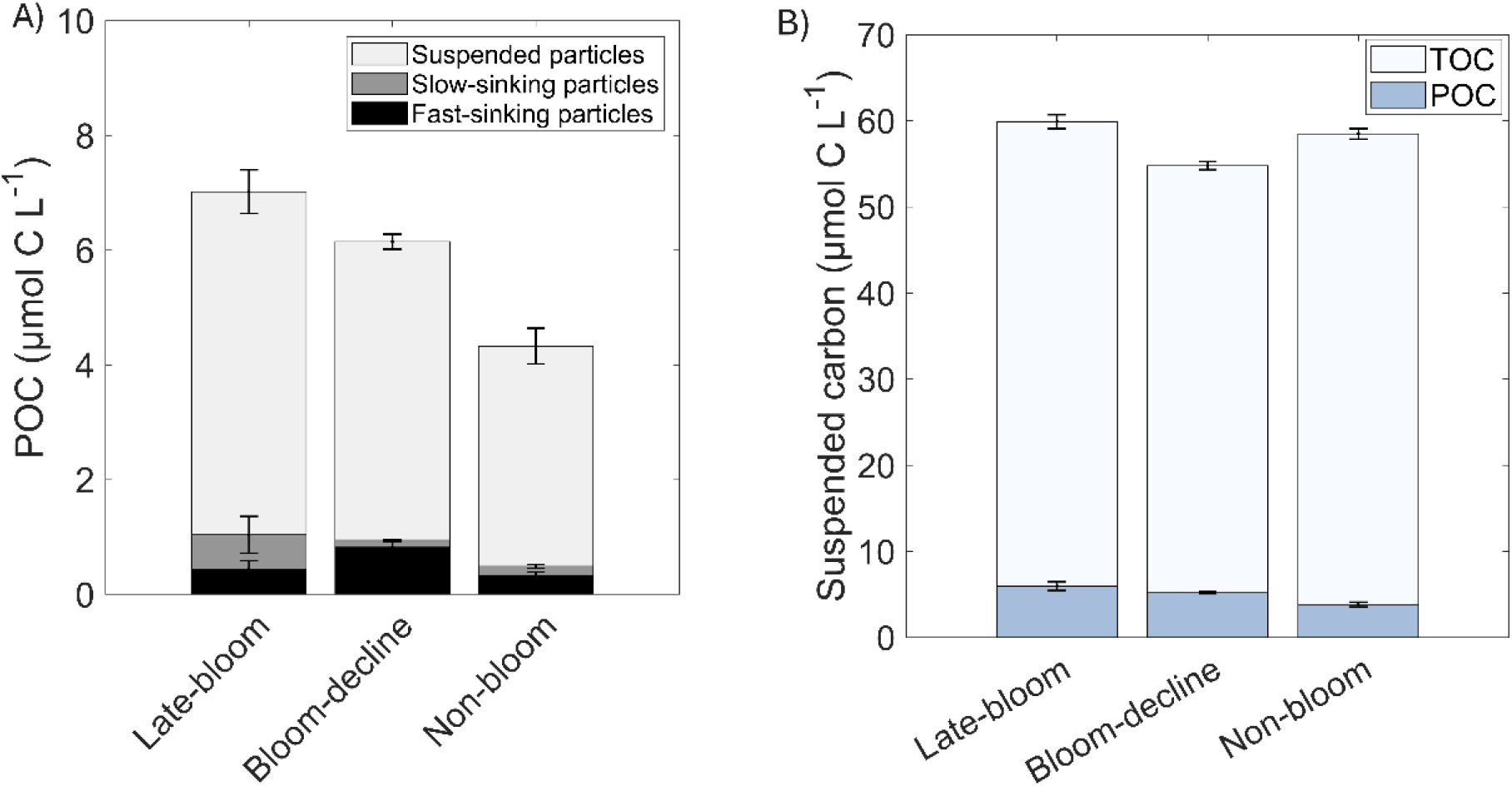
Initial concentrations of POC and TOC. **A)** Concentrations of POC (µmol C L^-1^) measured at the beginning of the incubation experiments (Day 0) in suspended particles (light grey), slow-sinking particles (dark grey), and fast-sinking particles (black). **B)** Comparison between initial concentrations of TOC (light) and POC (dark) in suspended particles. Error bars show propagated standard deviations from duplicate and triplicate measurements for POC and TOC, respectively.

**Figure 2.**
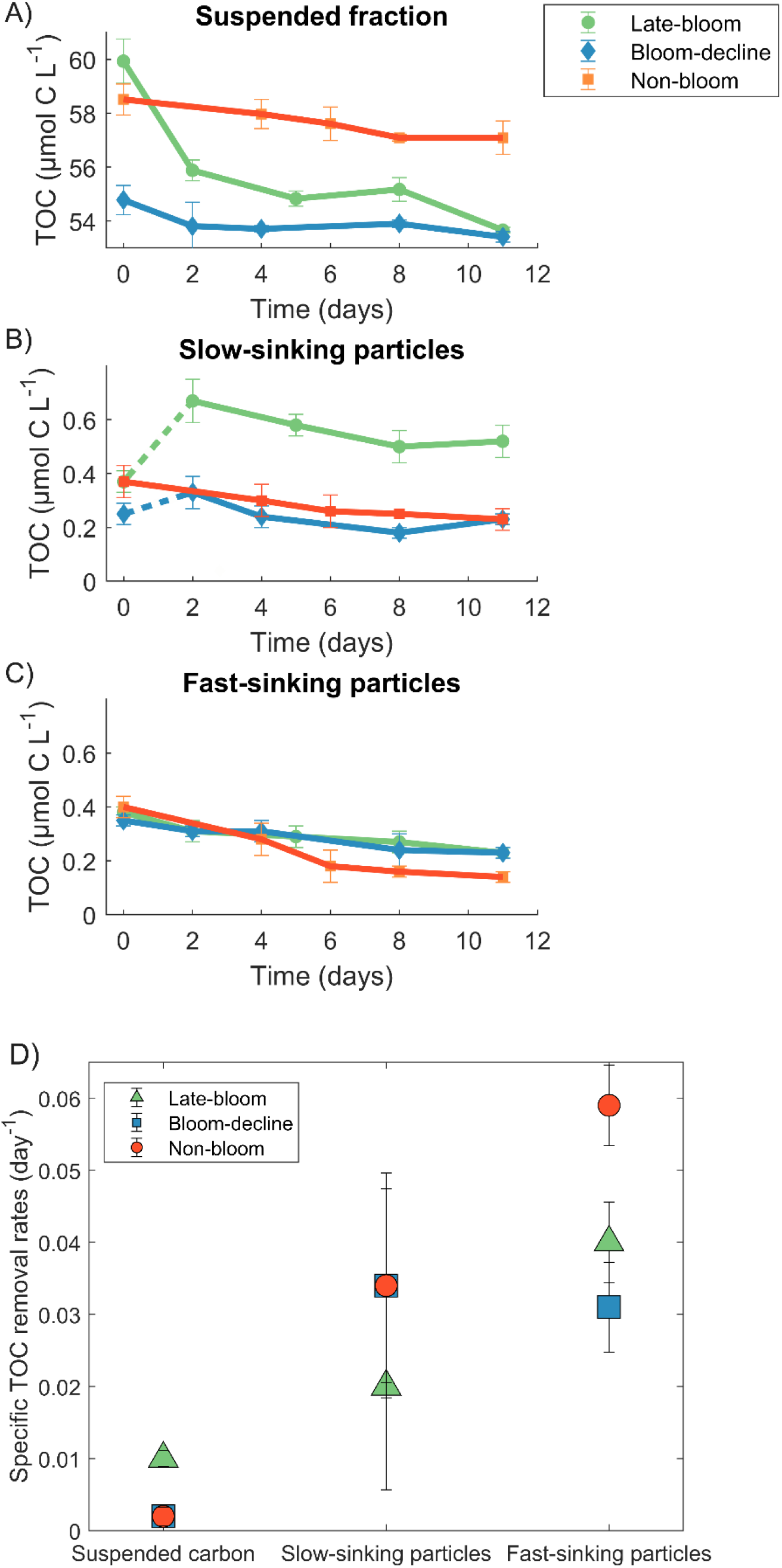
Temporal dynamics and removal of TOC over time across different stations and carbon fractions. **A, B, C)** Time series of TOC concentrations measured during the incubations at the late-bloom (green), bloom-decline (blue), and the non-bloom stations (orange). The data represent changes in TOC concentrations in **(A)** suspended carbon **(B)** in the slow-sinking and **(C)** fast-sinking particle fractions. Error bars indicate the standard deviation of the averages of triplicate samples propagated through the calculations. Dashed lines represent sections of the linear trend removed from rate calculations due to anomalously low values. **D)** Specific TOC removal rates (d^−1^) for suspended carbon, slow-sinking, and fast-sinking particles at three stations: late-bloom (green), bloom-decline (blue), and non-bloom (orange). Error bars show propagated standard deviations from triplicate measurements.

### Assessment of TOC removed by heterotrophic bacteria

The incubation experiments showed a statistically significant decline in TOC concentration over time across carbon fractions and stations (p ≤ 0.05), with three exceptions (**Figure 2A–C, Table S1**). Declines were not significant for slow-sinking particles at the late-bloom and bloom-decline stations, where the initial time point was omitted due to anomalously low values, or for suspended carbon at the bloom-decline station. TOC decline rates (µmol C L^−1^) over 11 days differed significantly between stations across all three carbon fractions, indicating that carbon turnover varied with bloom stage (**Table S1**).

We normalized TOC decline rates by initial TOC (specific removal rates, d^−1^) to account for differences in initial concentration and enable comparison across fractions and bloom stages (**Figure 2D**). At the late-bloom station, suspended carbon had ∼tenfold higher specific removal (0.01 d^−1^) than at the bloom-decline and non-bloom stations (0.002 d^−1^) (**Figure 2D**). This rate is underestimated because the decline was nonlinear; considering only the first two days gives 0.03 d^−1^ (steepest decline; **Figure 2A**). Specific removal rates of sinking particles were similar to suspended carbon at the late-bloom station but tenfold higher than at the other stations (0.02– 0.06 d^−1^; **Figure 2D**).

At all stations, specific removal rates increased substantially (4× to >10×) from suspended to sinking fractions. At the non-bloom and late-bloom stations, rates increased twofold from slow-to fast-sinking particles (**Figure 2D**). Slow-sinking particle removal rates at the late-bloom and bloom-decline stations should be interpreted with caution because the first data point was excluded (see methods).

### Characterization of exopolymer particles (TEP and CSP) in the water column

Whole water column POC at the late-bloom station was enriched in both CSP and TEP. Between 70 and 200 m, the CSP:POC and TEP:POC ratios were 2.6-fold and 2-fold higher, respectively, than at the bloom-decline station (**Figure 3**). Moreover, TEP:POC ratios consistently exceeded CSP:POC ratios, indicating a water column more enriched in TEP than in CSP. Microscopy revealed that *P. pouchetii* colonies, collected from the suspended fraction of the MSC at the late-bloom station, were beginning to disintegrate, but were still visibly containing *P. pouchetii* cells covered in CSP (**Figure 3**). CSP appeared to coat individual cells within colonies and free cells when colonies disintegrated. TEP was observed primarily as free-floating sheets not directly associated with *P. pouchetii* cells, although this material is known to form the mucus matrix of intact colonies (Mari et al., 2005) (**Figure 3**).

**Figure 3.**
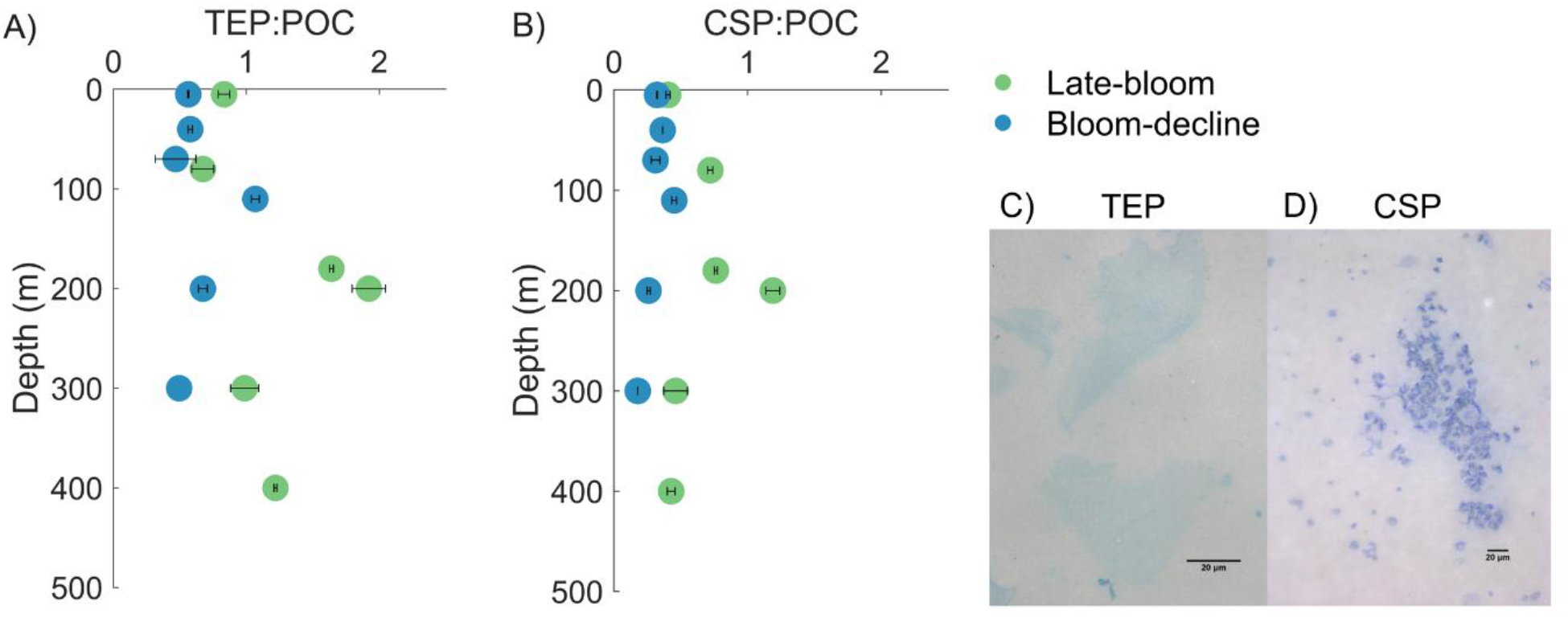
Exopolymer particle composition of suspended carbon at the late-bloom and bloom-decline stations. **A)** Vertical profiles of TEP:POC (µg Xeq µg^−1^ POC^−1^) and **B)** CSP:POC (µg BSAeq µg^−1^ POC^−1^) at the late-bloom (green) and bloom-decline (blue) stations. **C)** and **D)** show microscopy images from the suspended carbon fraction at a late-bloom station (station 6). **(C)** Suspended carbon was stained with Alcian Blue to visualize TEP, which appears not to be associated with *P. pouchetii* colonies; **(D)** Suspended organic carbon was stained with Coomassie Blue to visualize CSP, which are associated with *P. pouchetii* cells. Scale bars = 20 µm.

### Microbial community structure obtained using 16S rRNA gene metabarcoding

The microbial community structurediffered significantly between stations but not between MSC top, base and tray fractions. *P. pouchetii* dominated the photosynthetic community at all stations and MSC fractions, comprising >95% of the *chloroplast 16S rRNA gene (cp16S) ASV* counts in each MSC sample (**Figure S2**).

Across stations, the relative abundance of the *cp16S ASVs* dropped significantly relative to non-photosynthetic bacterial ASVs (**Figure 4A**). At the late-bloom station, the *cp16S* ASVs accounted for 53–58% of the total *16S* community, compared to 7-23% and 1-4.5% at the bloom-decline and non-bloom stations. *Thalassiosira* was the second most prevalent photosynthetic genus and occurred in slightly higher relative abundance at the non-bloom station, 0.9–4%, compared to 0.5-1.8% at the late-bloom and bloom-decline stations.

**Figure 4.**
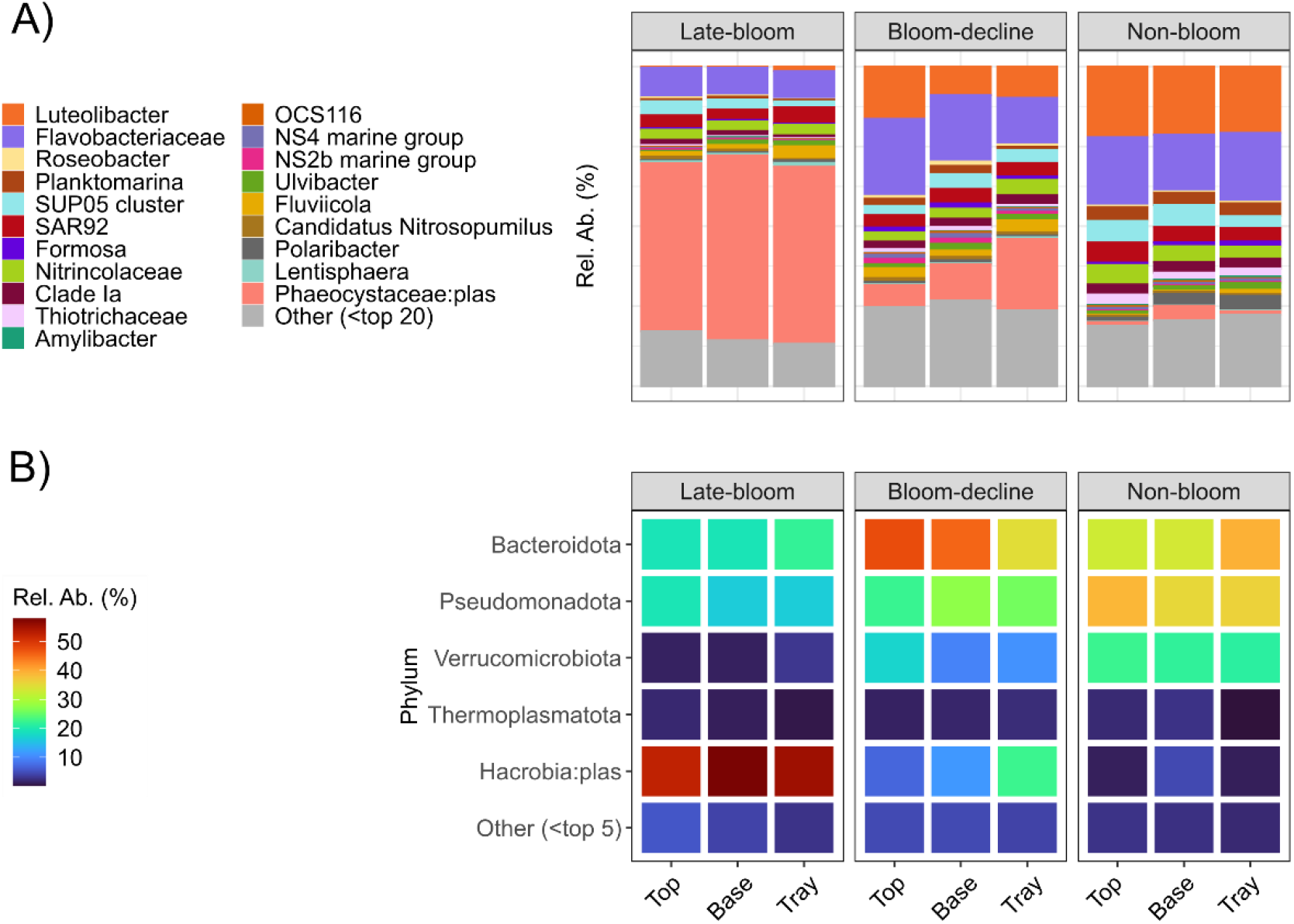
Relative abundances (Rel. Ab. (%)) of major taxa in the *16S* rRNA ASV analyses of microbial community observed in MSC top, base and tray fractions across stations. The “:plas” denotes that the taxonomic group is photosynthetic (i.e. Phaeocystaceae:plas). **(A)** Bar plot represents the relative abundance of the top 20 taxonomic groups by overall relative abundance at the highest possible taxonomic resolution. **(B)** Heatmap of the top 5 phyla.

The dominant prokaryotic phyla across stations were Bacteriodota, Pseudomonadota and Verrucomicrobia (**Figure 4B, Figure S3**). Notably, the relative abundance of Verrucomicrobia within the non-photosynthetic bacterial community increased markedly from 2.8–6.7% at the late-bloom station, to 11–19%, and 21–23% at the bloom-decline and non-bloom station, respectively (**Figure 4B, Figure S3**).

Beta diversity analysis, which compares the community structure between samples, revealed that microbial community composition was significantly influenced by station in both the entire *16S* community (R^2^ = 0.88, p = 0.001; **Figure 5A**) and the non-photosynthetic prokaryotic communities (excluding *cp16S*; R^2^ = 0.71, p = 0.001; **Figure 5B**). In contrast, the MSC top, base and tray carbon fractions did not differ significantly in microbial community structure (R^2^ = 0.05–0.13, p > 0.87).

**Figure 5.**
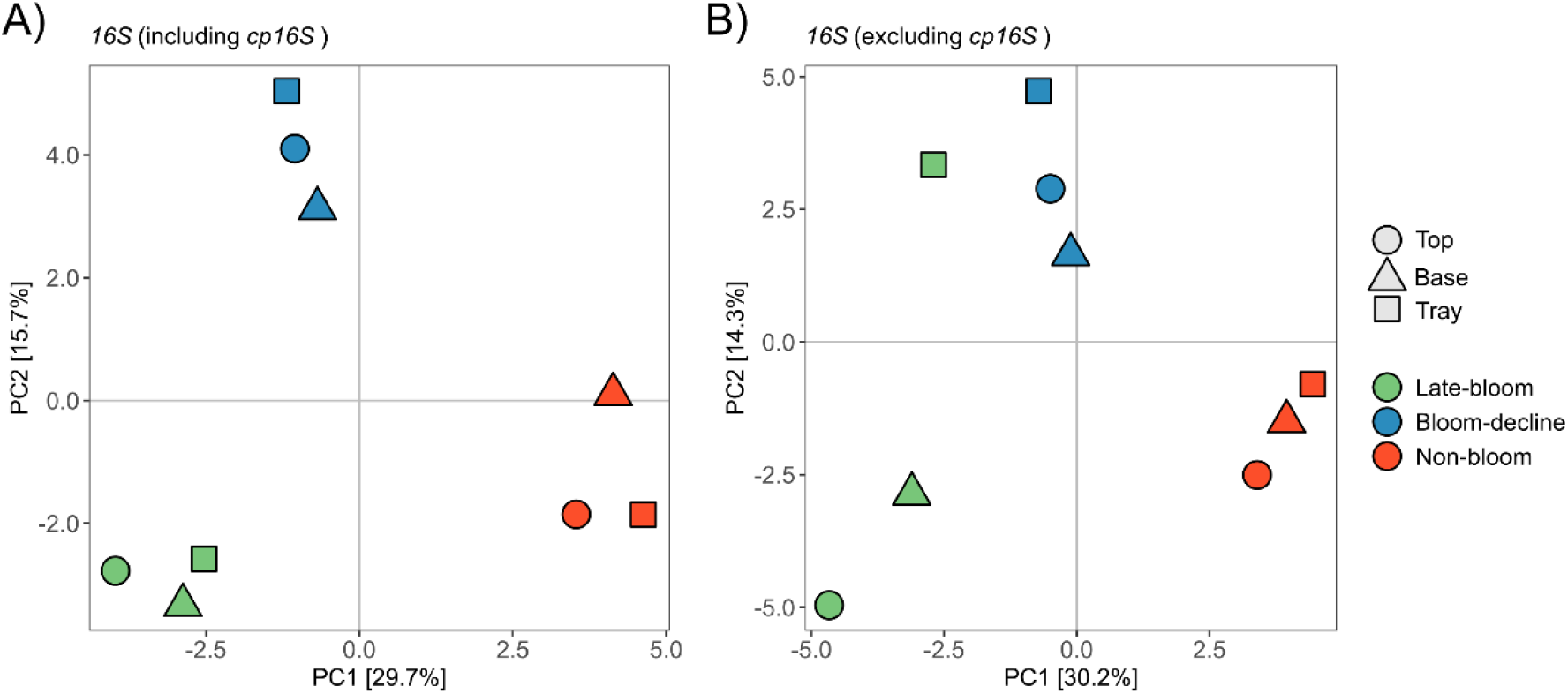
Principal component analyses (PCA) of microbial communities from MSC fractions. ASV data were CLR-transformed and plotted using PCA. Panel **(A)** represents the full *16S* community, and panel **(B)** represents the non-photosynthetic subset. In panels **(A)** and **(B)**, the MSC samples are colored by station, and their shape indicates the fractions. Community composition was significantly influenced by station in both the full **(A)**: R^2^ = 0.88, *p* = 0.001 and **(B)**: non-photosynthetic R^2^ = 0.71, *p* = 0.001 microbial communities, MSC fractions had no significant effect **(A, B)**: R^2^ = 0.05–0.13, *p* > 0.87.

The microbial community structure of the three carbon fractions cannot be considered fully independent, because the base fraction contains the suspended fraction, and the tray fraction encompasses both suspended and slow-sinking particles. Consequently, differences in community structure across carbon fractions may be obscured. We therefore applied indicator species analysis to identify ASVs within the non-photosynthetic bacterial community specifically associated with each carbon fraction. Seven indicator ASVs were found to be indicative of carbon fraction: ASVs in the Flavobacteriaceae, Spongiibacteraceae, Crocinitomicaceae, and Cyclobacteriaceae families for fast-sinking particles; two ASVs in Cryomorphaceae for slow-sinking particles; and an ASV in the SAR202 clade was indicative of the suspended carbon fraction (**Figure 6**). In addition, the indicator species analysis identified 95 ASVs within the non-photosynthetic prokaryotic community that were significantly associated with a given station. Of these, 27 ASVs were indicators of late-bloom, 13 ASVs were associated with the bloom-decline, and 55 ASVs were indicative of the non-bloom stage (**Figure S5**), suggesting ASV-specific differences driven by bloom stage. An ASV in the genus *Luteolibacter* (Verrucomicrobiota) was indicative of the non-bloom station (**Figure S4**). ASVs identified as indicators of the non-bloom station exhibited the greatest taxonomic diversity (**Figure S5**).

**Figure 6.**
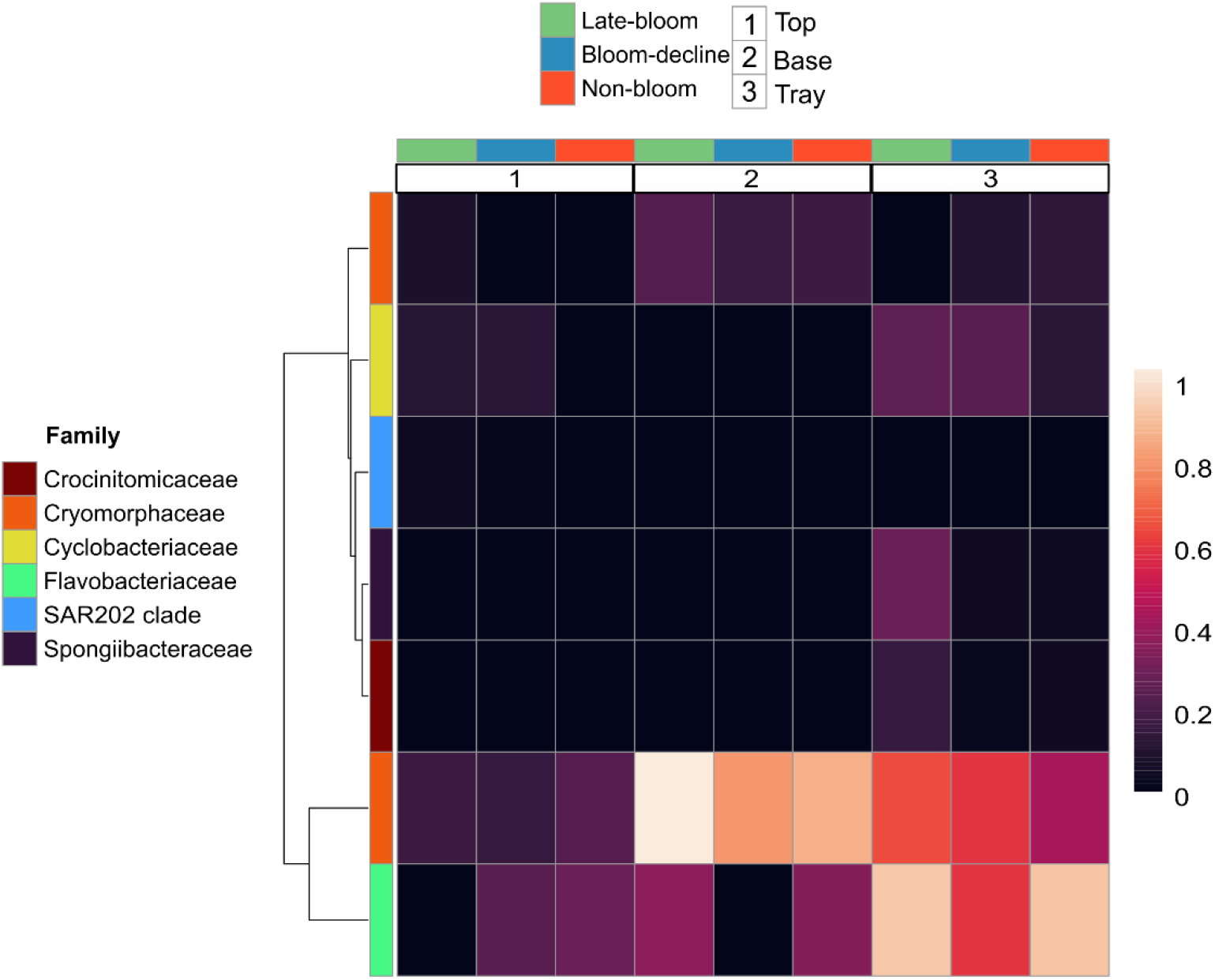
Relative abundance of ASV indicators associated with different carbon fractions within the non-photosynthetic bacterial *16S* V4V5 community at the Family level. The heatmap represents the relative abundance of each ASV in each MSC sample. The numbers indicate the MSC fraction, and the colors indicate the bloom stage in which the sample was collected.

## Discussion

### A novel approach to directly quantify TOC removed by microbial activity

Using a combination of MSC sampling and TOC incubation experiments, we directly quantified specific TOC removal rates for suspended carbon and sinking particle fractions. These rates, presumed to be driven by heterotrophic microbes, were measured at three stations representing different termination stages of a highly productive *P. pouchetii* bloom.

The measured specific TOC removal rates (0.002–0.06 d^−1^) fall within the global range of carbon-specific turnover rates inferred from oxygen consumption, reported as up to 1.8 d^−1^ for suspended carbon and 0.001–0.5 d^−1^ for sinking particles (Iversen & Ploug, 2013; Collins et al., 2015; McDonnell et al., 2015; Belcher et al., 2016; Garcia-Marti et al., 2021; Hemsley et al., 2023; Bressac et al., 2024, Petiteau et al. 2025). Rates measured for sinking particles were of the same order of magnitude as those reported for laboratory-made diatom aggregates (0.03 ± 0.01 d^−1^, Iversen & Ploug, 2013) and *in situ* sinking particles at similarly low temperatures (Bressac et al., 2024).

Our approach is the first to directly quantify TOC loss associated with microbial activity and concurrent microbial community responses on natural suspended and sinking carbon. In contrast, previous estimates derive turnover rates from oxygen consumption and require assumptions about respiratory quotients, which can vary and introduce uncertainty (Robinson, 2005; Stephens et al., 2025). By incubating unfiltered samples and measuring TOC directly, our rates integrate particulate solubilization by particle-attached bacteria, subsequent DOC production, and DOC removal by free-living bacteria. Subtracting MSC top concentrations from base concentrations (base–top) isolates slow-sinking particles, while subtracting base from tray concentrations (tray–base) isolates fast-sinking particles. This approach corrects for background DOC present at the time of sampling while retaining DOC produced during particle solubilization, yielding conservative yet direct estimates of carbon removal for each carbon fraction. Additional methodological considerations are provided in the Supplementary Material. In the following sections, we place these specific removal rates in an oceanographic context to evaluate how bloom-stage–dependent particle lability influences microbial assembly and processing of organic carbon.

### Late-bloom exopolymer enrichment drives rapid microbial degradation of suspended P. pouchetii carbon

At the late-bloom station, suspended *P. pouchetii* carbon was rapidly removed, with most TOC decline occurring within the first two days, consistent with an immediate microbial response (**Figure 2**). This elevated turnover coincided with pronounced enrichment of TEP and CSP in the water column POC (**Figure 3**), indicating that carbon composition, rather than POC standing stock (**Figure 1A**), best explains the observed differences in removal rates of suspended carbon across stations. TEP likely formed abiotically through coagulation of dissolved precursors exuded by abundant fresh colonies and released by decaying porous colonies (Mari et al., 2005). Whereas CSP was primarily associated with growing cells or cells released from decaying colonies. The concurrent release of TEP and CSP provided labile polysaccharides and proteins, which are known to trigger rapid microbial activity (Passow, 2002; Koch et al., 2014; Busch et al., 2017). Multivariate analysis linked variation in heterotrophic prokaryotic community structure with bloom stages, consistent with bloom-driven changes in particle lability. Previous incubation work with concentrated *Phaeocystis* mucus reported stable bacterial community composition during rapid substrate loss (∼50% TOC loss over 11 days at 12 °C; k ∼ 0.063 d^−1^; temperature-adjusted k ∼ 0.036 d^−1^ at 4 °C), with community shifts emerging only after most labile substrate was consumed (Janse et al., 1999; Janse et al., 2000). These results are consistent with our observation of intense early degradation followed by longer-term community restructuring. Collectively, the data indicates that exopolymer-rich carbon during late blooms enhances microbial remineralization of *Phaeocystis* colonies below the mixed layer, likely reducing their export efficiency.

### Microbial mechanisms driving limited export of Phaeocystis-derived organic carbon

Enhanced microbial breakdown of TEP and CSP-rich suspended carbon is shown to reduce carbon transfer to depth. A TOC removal rate of 0.03 d^−1^ (steepest decline; **Figure 2A**) corresponds to a half-life of ∼23 days at 4 °C, implying that a large fraction of this labile carbon may be recycled before substantial sinking. Consistently, organic carbon export flux measured at the base of the primary production zone during the expedition did not differ significantly between the late-bloom (station 9; 6.2 ± 2.9 mmol C m^−2^ d^−1^) and bloom-decline stations (station 28; 5.1 ± 2.9 mmol C m^−2^ d^−1^, Roca-Martí et al., 2025) supporting limited vertical export of *P. pouchetii*-derived carbon during the late-bloom stage.

Although particle sinking velocities were not directly measured, the buoyant nature of TEP can reduce particle sinking, prolong residence in the upper water column, and increase exposure to bacteria, releasing DOC at shallow depths and further limiting vertical export depths (Romanelli et al., 2023; Chajwa et al., 2024; Alcolombri et al., 2024). Enhanced lability could potentially stimulate zooplankton grazing. Although grazing rates were not measured during the expedition, integrated zooplankton biomass (100–150 m) was similar between the late-bloom and bloom-decline stations and lower at the late-bloom station than at the non-bloom station (McGinty et al., *pers. comm*.). This observation suggests low grazing pressure on *Phaeocystis* colonies at the late-bloom stage, which is consistent with previous investigations (Saiz et al., 2013).

Together, these observations indicate that rapid microbial recycling of CSP- and TEP-rich suspended carbon is associated with low export efficiency during late-stage *P. pouchetii* blooms.

### Contrasting removal dynamics and bacteria community assembly of suspended and sinking organic carbon across bloom stages

Our results show that suspended and sinking organic carbon processing varies with bloom stage. At the late-bloom station, TOC removal rates of fast-sinking particles were of the same order of magnitude as suspended carbon (**Figure 2B**), indicating comparable lability. This reflects a common origin from similarly aged *P. pouchetii* colonies and is supported by Particulate Hydrolyzable Amino Acids (PHAA) degradation indices and labile amino acid concentrations, which did not differ between fractions (Cisternas-Novoa *pers. comm*.). These findings suggest that exopolymer-rich suspended material can act as microbial hotspots comparable to sinking particles, with TEP- and CSP-enriched carbon during late-stage *P. pouchetii* blooms sufficiently labile to sustain high microbial turnover, providing a potential pathway for carbon processing in the upper mesopelagic (Hemsley et al., 2023; Burd et al., 2010). In contrast, at bloom-decline and non-bloom stations, removal rates of sinking particles exceeded suspended carbon by roughly an order of magnitude (**Figure 2B**), indicating that suspended matter had become largely refractory. This pattern reflects advanced bloom decay, with fewer intact *P. pouchetii* colonies, more detritus, and elevated degradation indices (phaeopigment-to-Chl a, Laget et al., 2025). Carbon fractions at these stages were enriched in Verrucomicrobiota, primarily *Luteolibacter*, reaching ∼21–23% of total abundance at the non-bloom station, consistent with their specialization in degrading complex polysaccharides (Sichert et al., 2020) and previous observations during the decline of *Phaeocystis* and diatom blooms (Orellana et al., 2022; Shi et al., 2023; Wilkie & Orellana, 2024; Skouroliakou et al., 2024).

Microbial community structure mirrored bloom-stage–dependent carbon lability. Bloom-decline and non-bloom stages exhibited higher bacterial α-diversity than late-bloom (**Figure S6**) and increased non-photosynthetic bacteria relative to photosynthetic taxa (**Figure 4**), consistent with observations that active blooms support low-diversity bacterial communities dominated by opportunistic lineages, while post-bloom conditions promote niche diversification as organic matter becomes more heterogeneous and chemically complex (Buchan et al., 2014; Wemheuer et al., 2014). Additional diatom-derived organic matter at the non-bloom station likely enhanced substrate diversity for heterotrophic metabolism. Fast-sinking particles at this station had the highest specific TOC removal rate (k ∼ 0.06 d^−1^; half-life ∼11.6 days), suggesting a highly labile component, likely diatom-derived. UVP-derived flux profiles corroborate this, showing strong attenuation of organic carbon export (Laget et al., 2025).

### Implications for the assessments of organic carbon sequestration in the biological carbon pump

Bloom-stage–dependent lability differences among carbon fractions may have important implications for ocean carbon sequestration. Across bloom stages, specific TOC removal rates of slow- and fast-sinking particles were two-to >10-fold higher than for suspended carbon (**Figure 2D**). Fast-sinking particles also had roughly twofold higher rates than slow-sinking particles at two of the three stations, suggesting that organic carbon degradability increases from suspended to fast-sinking fractions.

Vertical remineralization profiles are influenced not only by remineralization rates but also by particle sinking velocity. Fast-sinking particles are often assumed to transport carbon deeper into the mesopelagic, while slow-sinking particles are remineralized at shallower depths, assuming similar remineralization rates (Buesseler et al., 2009). Here, fast-sinking particles exhibited elevated remineralization rates compared to slow-sinking particles and suspended particles, reducing differences in their remineralization length scales by roughly a factor of two. Together with recent MSC studies (Baker et al., 2017), these results suggest that slowly sinking particles—previously under-sampled— may contribute substantially to organic carbon sequestration and mesopelagic microbial metabolism (Burd et al., 2010). Broader observations across particle fractions and environmental conditions are needed to confirm these biogeochemical impacts.

Many Earth System Models (ESMs) represent sinking fluxes using fast- and slow-sinking particle pools with fixed remineralization rates (e.g. ROMS-BEC, PISCES-v2, FESOM2.1– REcoM3) (Gruber et al., 2006; Aumont et al., 2015; Gürses et al., 2023). However, particle-specific remineralization rates differ between models: PISCES-v2 and FESOM2.1–REcoM3 assume the same rates for both pools, while ROMS-BEC assumes rates three times higher for slow-sinking particles. Our assessment is not exhaustive because our dataset is limited, however it highlights that these assumptions may overlook differential microbial processing and underscores the need to revise ESM parameterizations to better reflect particle-associated microbial activity.

## Conclusions

Our results demonstrate that bloom-stage–dependent shifts in particle lability influence microbial assembly and processing of organic carbon and its fate in the upper mesopelagic. During the late stage of *P. pouchetii* blooms, suspended, exopolymer-rich carbon can contribute to microbial remineralization rates similar to that of sinking particles, providing an explanation for the limited export of *P. Pouchetii*-derived carbon into the deep ocean. Bloom progression also modulates prokaryotic community composition. While overall microbial composition differed little between carbon fractions, Flavobacteriaceae, Spongiibacteraceae, Crocinitomicaceae, and Cyclobacteriaceae families were indicative of fast-sinking particles, suggesting selective colonization of rapidly exported material. With *Phaeocystis* blooms increasingly replacing diatom-dominated blooms in subpolar regions (Devred et al., 2025), understanding stage-specific carbon dynamics and microbial processing is essential for accurately predicting carbon fluxes. Future work should aim to quantify how exopolymers produced at different bloom stages regulate bacterial activity and alter particle buoyancy and export efficiency at the single-particle level.

Overall, we find that TOC removal rates of slow- and fast-sinking particles were greater than for suspended carbon and that fast-sinking particles exhibited faster TOC removal rates than the slow-sinking particles. This finding likely has important implications for the assessments of the importance of slow-sinking particles in global carbon storage and for sustaining mesopelagic communities.

## Acknowledgements

E.R. and D.A.S. were supported by NASA Grant 80NSSC17K0692. R.S.G. was supported by the NSERC Canada Graduate Scholarship-Doctoral Award with the Fisheries and Oceans Canada Aquatic Science Supplement provided by DFO, as well as the Nova Scotia Graduate Scholarship from Dalhousie University. CCN was supported by Ocean Frontier Institute’s research project “The Northwest Atlantic Biological Carbon 598 Pump” (NWA-BCP). CAC was supported by Simons Foundation International’s BIOS-SCOPE project. UP and JLR were supported by the Canada Research Chair Program. This study is part of the Ocean Frontier Institute’s research project “The Northwest Atlantic Biological Carbon 598 Pump” (NWA-BCP). We are sincerely grateful to the crew and scientific party aboard the RV Celtic Explorer 599 during the BELAS-1 expedition and the whole NWA-BCP team. We also thank the Hervé Claustre group (CNRS, Laboratoire d’Océanographie de Villefranche, France) for the collection, extraction, and HPLC analysis of CTD pigment samples.

## Data accessibility statement

All raw sequence data and analysis code are publicly available (https://github.com/BeccaStevens-Green/MSC_samples_in_the_LS_BELAS-1_TOC_rem/tree/main).

