## Supplemental material for "Particle lability drives degradation dynamics and bacterial community assembly during a *Phaeocystis* bloom decline"

*Further methodological considerations*

To our knowledge, this is the first time that TOC removal rates have been directly resolved for both slow-sinking and fast-sinking particles. The precision of the TOC analyzer at these concentrations is limited (±0.7 µM C), making it challenging to detect a signal specifically attributable to sinking particles. By applying the MSC methodology, we were able to concentrate slow- and fast-sinking particles, thereby improving the signal-to-noise ratio and allowing us to confidently resolve changes in TOC. The MSC approach has previously been used to estimate microbial activity on different particle fractions (Garcia-Martí et al., 2021; Belcher et al., 2016; Hemsley et al. 2023), although it does not allow for direct quantification of microbial activity uniquely associated with sinking particles (Garcia-Martí et al., 2021). As described in Section 2.1, the MSC base fraction contains both suspended carbon (particulate and dissolved) and slow-sinking particles, whereas the tray fraction contains suspended carbon and slow-sinking and fast-sinking particles. Based on these considerations, we cannot exclude the possibility that part of the activity attributed to sinking particles reflects the activity of free-living bacteria consuming dissolved organic carbon that was either present at the start of the incubations or released by particle-attached bacteria during remineralization. Consequently, our experiments capture not only the remineralization and solubilization processes carried out by particle-associated bacteria but also, at least partially, the activity of free-living bacteria. Importantly, sinking particles in the ocean are never found in isolation: they are embedded in a matrix of suspended particles and surrounded by dissolved organic matter. Thus, it is ecologically realistic that free-living bacteria benefit from the dissolved organic carbon released during particle remineralization (Cho & Azam, 1988; Kiørboe & Jackson, 2001; Collins et al., 2015; Garcia-Martí et al., 2021).

*Identification of most abundant cp16S ASVs in the dataset*

There was a total of 16 *cp16S* ASVs present in the 9 samples. The dominant ASV which contributed 97.7% of the community is part of the Hacrobia phyla and was further classified as *Phaeocystis pouchetii* using full *18S* rRNA gene sequencing. The other 3 ASVs in Hacrobia were classified as *Teleaulax* (0.30% of *cp16S*), another *Phaeocystis* (0.014% of *cp16S*) and one in the Prymnesiophyceae family (0.0033% of *cp16S*). The top 2/7 ASVs in the Stramenopiles family were classified as belonging to *Thalassiosira sp.*(1.65%) and the Chrysophyceae family (0.22%).

**Rerefences**

García-Martín, E.E., Davidson, K., Davis, C.E., Mahaffey, C., Mcneill, S., Purdie, D.A. and Robinson, C., 2021. Low contribution of the fast-sinking particle fraction to total plankton metabolism in a temperate shelf sea. *Global Biogeochemical Cycles*, *35*(9), p.e2021GB007015.

Belcher, A., Iversen, M., Giering, S., Riou, V., Henson, S.A., Berline, L., Guilloux, L. and Sanders, R., 2016. Depth-resolved particle-associated microbial respiration in the northeast Atlantic. *Biogeosciences*, *13*(17), pp.4927-4943.

Hemsley, V., Füssel, J., Duret, M.T., Rayne, R.R., Iversen, M.H., Henson, S.A., Sanders, R., Lam, P. and Trimmer, M., 2023. Suspended particles are hotspots of microbial remineralization in the ocean's twilight zone. *Deep Sea Research Part II: Topical Studies in Oceanography*, *212*, p.105339.

Cho, B.C. and Azam, F., 1988. Major role of bacteria in biogeochemical fluxes in the ocean's interior. *Nature*, *332*(6163), pp.441-443.

Kiørboe, T. and Jackson, G.A., 2001. Marine snow, organic solute plumes, and optimal chemosensory behavior of bacteria. *Limnology and Oceanography*, *46*(6), pp.1309-1318.

Collins, J.R., Edwards, B.R., Thamatrakoln, K., Ossolinski, J.E., DiTullio, G.R., Bidle, K.D., Doney, S.C. and Van Mooy, B.A., 2015. The multiple fates of sinking particles in the North Atlantic Ocean. *Global Biogeochemical Cycles*, *29*(9), pp.1471-1494.

**Supplemental Tables**

**Suspended carbon:**

**A) Weighted linear regression**

| Group | Slope (µmol C $L^{-1}$ $d^{-1}$) | p-value |
|---|---|---|
| Late bloom | -0.2853 | 0.0422 |
| Bloom decline | -0.0352 | 0.4567 |
| Non bloom | -0.1699 | 0.0133 |

**B) ANCOVA (analysis of covariance)**

| Source | SumSq | DF | MeanSq | F | pValue |
|---|---|---|---|---|---|
| Total | 814.93 | 14 | 58.209 | | |
| Model | 786.47 | 5 | 157.29 | 49.743 | 2.754e-06 |
| Linear | 751.64 | 3 | 250.55 | 79.234 | 8.62e-07 |
| Nonlinear | 34.832 | 2 | 17.416 | 5.508 | 0.0274 |
| Residual | 28.459 | 9 | 3.1621 | | |

**Slow-sinking particles:**

**C) Weighted linear regression**

| Group | Slope (µmol C $L^{-1}$ $d^{-1}$) | p-value |
|---|---|---|
| Late bloom | −0.0158 | 0.1308 |
| Bloom decline | −0.0035 | 0.7489 |
| Non-bloom | −0.0126 | 0.0086 |

**D) ANCOVA (analysis of covariance)**

| Source | SumSq | DF | MeanSq | F | pValue |
|---|---|---|---|---|---|
| Total | 151.66 | 12 | 12.638 | | |
| Model | 143.22 | 5 | 28.643 | 23.759 | 0.00029518 |
| Linear | 141.12 | 3 | 47.041 | 39.020 | 9.714e−05 |
| Nonlinear | 2.0935 | 2 | 1.0468 | 0.86827 | 0.46042 |
| Residual | 8.4389 | 7 | 1.2056 | | |

**Fast-sinking particles:**

**E) Weighted linear regression**

| Incubation | Slope (µmol C $L^{-1}$ $d^{-1}$) | p-value |
|---|---|---|
| Late bloom | −0.0131 | 0.0034 |
| Bloom decline | −0.0104 | 0.0025 |
| Non-bloom | −0.0223 | 0.0167 |

**F) ANCOVA (analysis of covariance)**

| Source | SumSq | DF | MeanSq | F | pValue |
|---|---|---|---|---|---|
| Total | 145.51 | 14 | 10.393 | | |
| Model | 139.43 | 5 | 27.886 | 41.308 | 6.1012e−06 |
| Linear | 132.50 | 3 | 44.168 | 65.426 | 1.9652e−06 |
| Nonlinear | 6.9283 | 2 | 3.4641 | 5.1314 | 0.032572 |
| Residual | 6.0758 | 9 | 0.67508 | | |

**Table S1**: **A, C, E)** Weighted linear regression results for suspended carbon, slow-sinking and fast-sinking particles: Temporal trends in TOC concentrations for each carbon fraction and station were quantified using weighted linear regression of mean TOC values (µmol C $L^{-1}$) across the 11 incubation days, with weights equal to the inverse variance of the three technical replicates. Slopes and associated p-values indicate the significance of TOC changes over time. Significant negative slopes indicate a measurable decrease in TOC. **B,D,F)** ANCOVA results for suspended carbon, slow-sinking and fast-sinking particles: Analysis of covariance (ANCOVA) tested whether temporal slopes differed among stations within each carbon fraction, using time as a covariate and the group × time interaction to assess slope differences. The table reports sum of squares (SumSq), degrees of freedom (DF), mean squares (MeanSq), F-values, and p-values for the model, linear component, nonlinear component, and residuals. A $p < 0.05$ indicates that the rates of TOC decline differ significantly among stations within each carbon fraction.

### Supplemental Figures

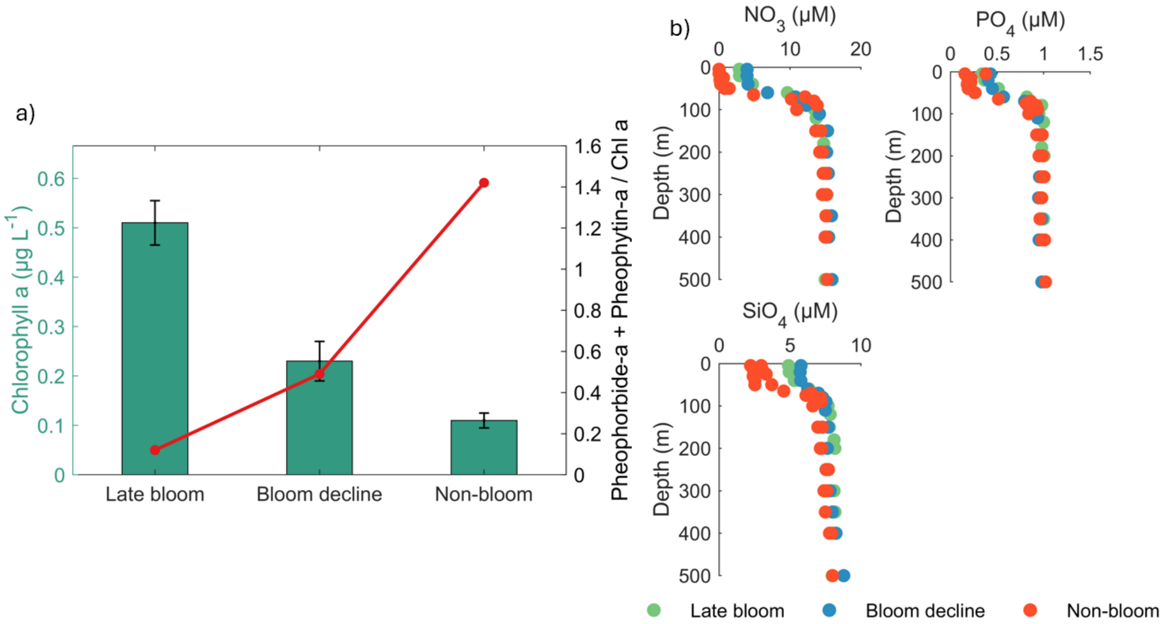


**Figure S1**. **(a)** Water column chlorophyll-a (Chl-a) concentrations measured by the fluorescence sensor on the CTD, which was deployed at the same depth prior to the MSC deployment (green bars) and the ratio of pheophorbide-a plus pheophytin-a to Chl-a, a biomarker for organic matter degradation in the water column, calculated from HPLC pigment measurements taken from the CTD samples collected during the same deployment (red line). **(b)** Profiles of the concentration of nitrate ($NO_3$), phosphate ($PO_4$), silicic acid ($SiO_4$) measured on water collected with Niskin bottles at the late bloom station (green), bloom decline station (blue) and the non-bloom station (orange) on the same days of the MSC deployments.

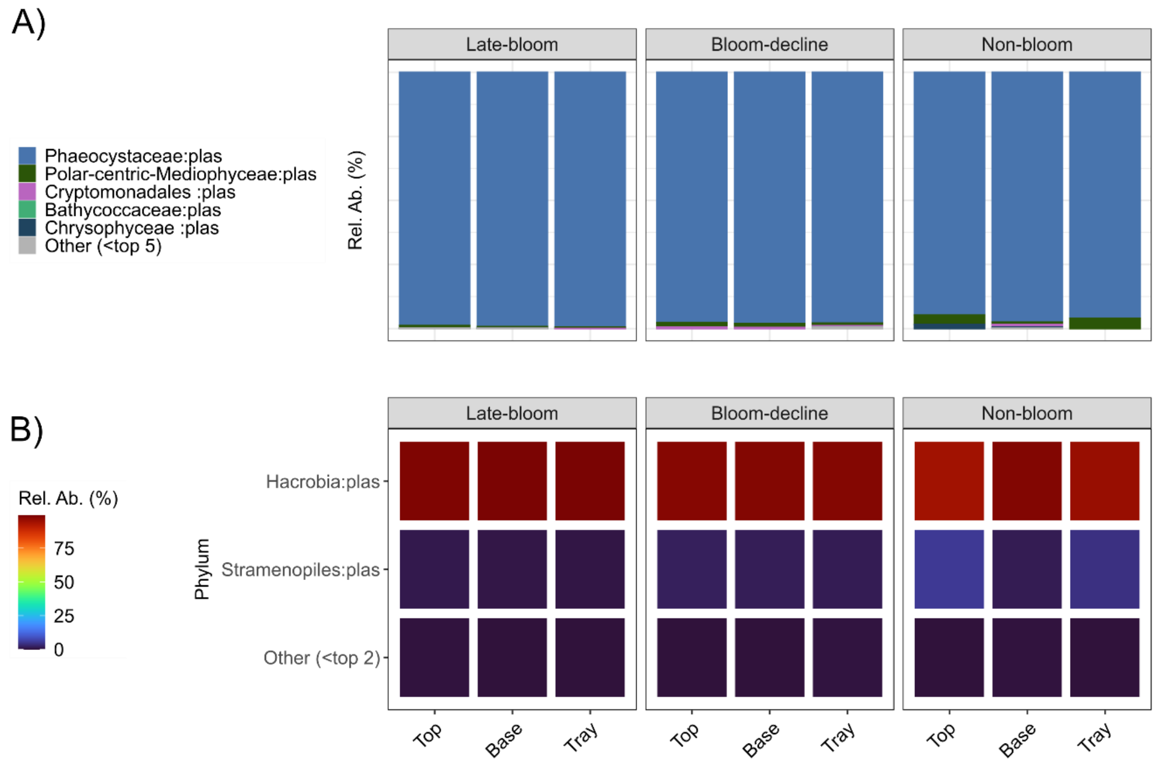


**Figure S2.** Relative abundances (Rel. Ab. (%)) of major photosynthetic taxa (*cp16S*) observed in MSC top, base and tray fractions across stations. **(A)** Bar plot represents the relative abundance of the top 5 taxonomic groups. ASVs are annotated to the most resolved level of taxonomic annotation. **(B)** Heatmap of the top 2 photosynthetic phyla including Hacrobia which is dominated by *Phaeocystis pouchetii* (average = 97.7%) and the Stramenopiles including mainly centric diatoms such as *Thalassiosira sp*.

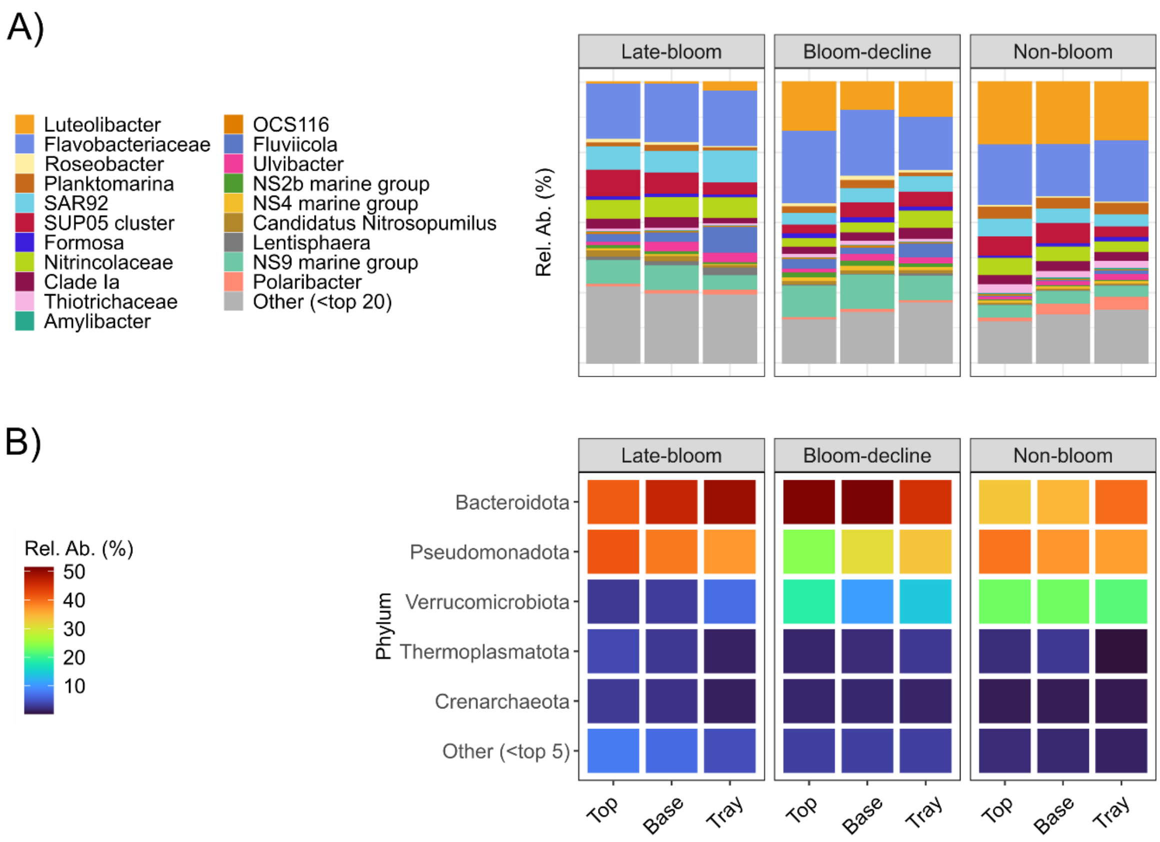


**Figure S3.** Relative abundances (Rel. Ab. (%)) of major taxa in the non-photosynthetic *16S* rRNA microbial community observed in MSC top, base and trays fractions across stations. **(A)** Bar plot represents the relative abundance of the top 20 taxonomic groups by overall relative abundance at the highest possible taxonomic resolution. **(B)** Heatmap of the top 5 phyla.

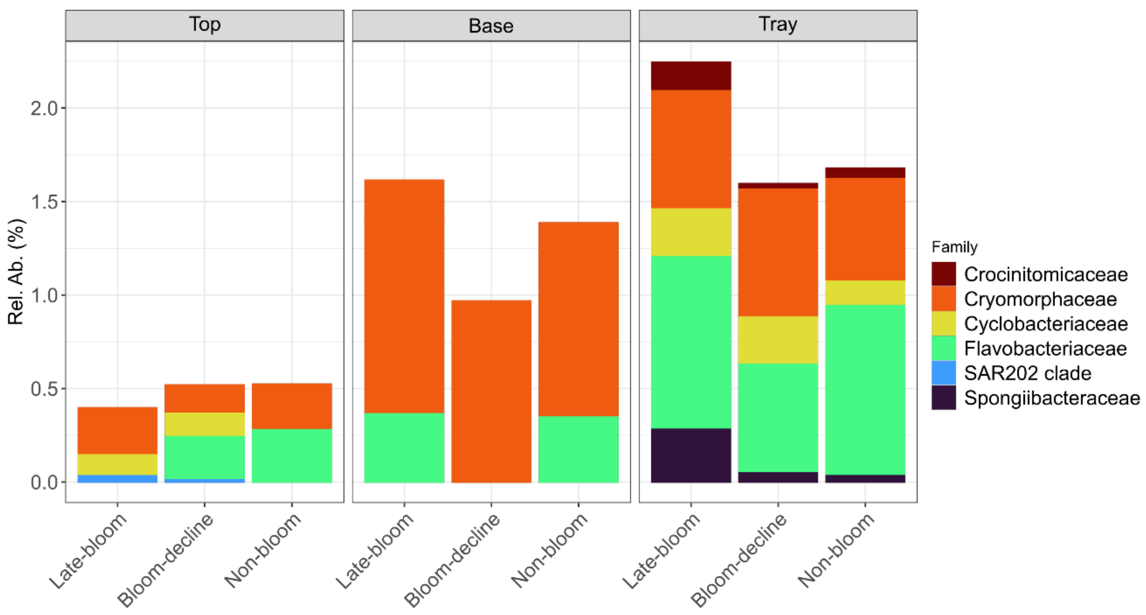


**Figure S4.** Relative abundance of ASV indicators associated with different carbon fractions within the non-photosynthetic bacterial ASV pool classified at the Family level. Indicator ASVs from each bloom stage are plotted in the same relative abundance plot to highlight the shifts in community composition of ASVs indicative of MSC carbon fractions.

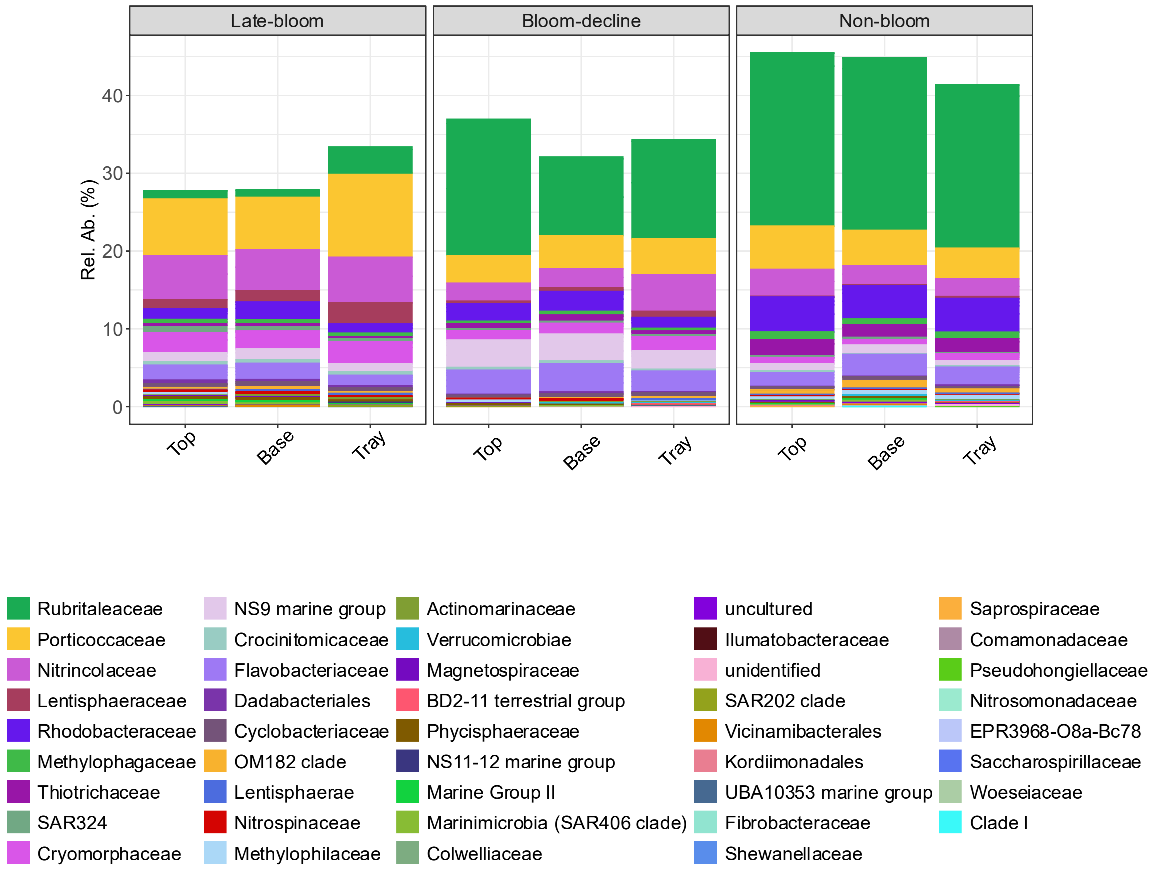


**Figure S5:** Relative abundance of the ASV indicators of bloom condition (late-bloom, bloom-decline and non-bloom) determined from the non-photosynthetic bacterial ASV pools. The colors represent distinct taxonomic groups at the Family level. Indicator ASVs from each bloom stage are plotted in the same relative abundance plot to highlight the shifts in community composition of indicator ASVs throughout the bloom progression.

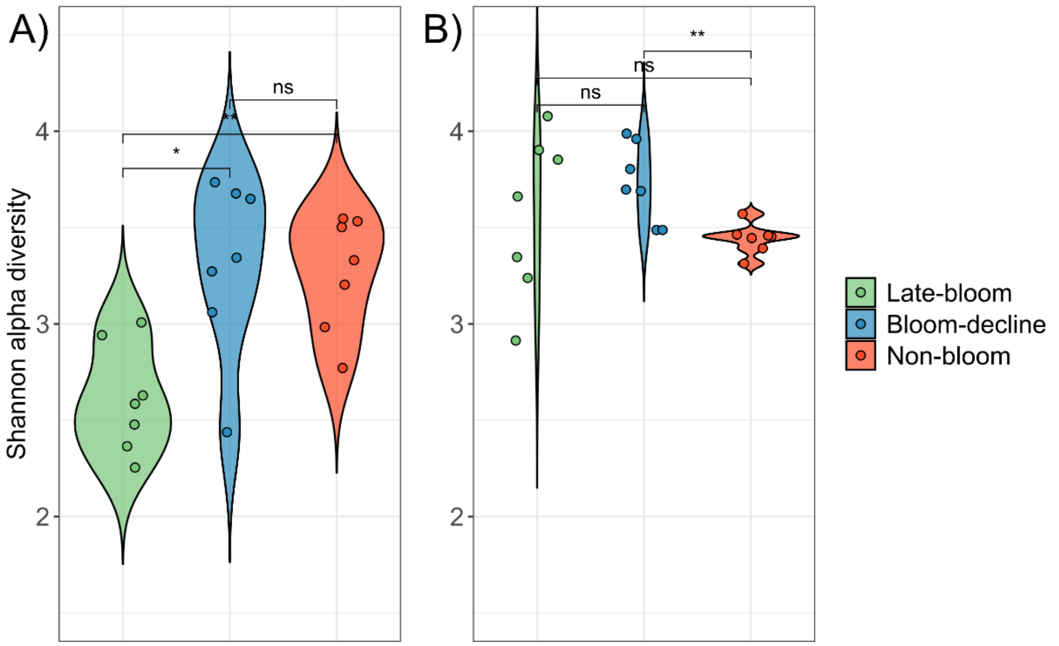


**Figure S6.** Violin plots of Shannon alpha diversity from MSCs and associated water column samples. Panels **(A)** and **(B)** show Shannon alpha diversity of the full microbial community (including *cp16S*, rarefied to 1,450 reads) and the non-photosynthetic bacterial subset (excluding *cp16S*, rarefied to 1,300 reads), respectively. Samples are grouped by bloom stage and significance is assessed using Kruskal-Wallis and pairwise Wilcoxon tests (Benjamini-Hochberg corrected). Points represent individual samples, and colors correspond to bloom stages. Asterisks indicate significant pairwise differences (* $p < 0.05$, ** $p < 0.01$), and "ns" indicates non-significant comparisons.
